# Sex- and Age-dependent Skin Mechanics – A Detailed Look in Mice

**DOI:** 10.1101/2023.03.08.531781

**Authors:** Chien-Yu Lin, Gabriella P. Sugerman, Sotirios Kakaletsis, William D. Meador, Adrian T. Buganza, Manuel K. Rausch

## Abstract

Skin aging is of immense societal and, thus, scientific interest. Because mechanics play a critical role in skin’s function, a plethora of studies have investigated age-induced changes in skin mechanics. Nonetheless, much remains to be learned about the mechanics of aging skin. This is especially true when considering sex as a biological variable. In our work, we set out to answer some of these questions using mice as a model system. Specifically, we combined mechanical testing, histology, collagen assays, and two-photon microscopy to identify age- and sex-dependent changes in skin mechanics and to relate them to structural, microstructural, and compositional factors. Our work revealed that skin stiffness, thickness, and collagen content all decreased with age and were sex dependent. Interestingly, sex differences in stiffness were age induced. We hope our findings not only further our fundamental understanding of skin aging but also highlight both age and sex as important variables when conducting studies on skin mechanics.

## INTRODUCTION

Skin ages. This inevitability is of significant societal, commercial, and thus scientific interest. Because of skin’s exposed role, it ages both intrinsically and extrinsically^1^. The former is driven by senescent mechanisms, while the latter is primarily driven by environmental exposures such as UV light^2^. The complex interplay between both intrinsic and extrinsic mechanisms sets skin apart from other organs and makes its aging a complex phenomenon that remains incompletely characterized. This is especially true once sex dependence is considered.

Naturally, aging also leads to biomechanical changes in skin^3^. Age-induced alterations in collagen and elastin quality and quantity change skin’s ability to deform under load and to resist injury^4–6^. Motivated by skin’s clinical and aesthetic importance, age-dependent alterations in skin mechanics have been subject to tremendous scientific curiosity^7^. Notwithstanding these efforts, there is little to no consensus on most aspects of age-induced biomechanical changes in skin; let alone when sex differences are considered.

Disagreements include fundamental questions such as whether skin stiffens^8–11^ with age or softens^12^, whether skin thins with age or not^13–15^, whether collagen content decreases with age^16^ or not^17,18^, and whether skin is stiffer in women^11^ or doesn’t differ^19,20^. Reasons for such dissonance are multi-fold and include: i) Methodological differences, i.e., drastically differing means to quantify mechanical changes, which include *in-vivo* suction^9,21,22^, torsion^8^, optical methods^23^, compression^24^, indentation^25,26^, and post-mortem test methods such as uniaxial extension^3,27–29^. ii) Skin biomechanical properties depend on location^30–32^ as skin function varies across our bodies while different degrees of UV exposure lead to locally-varying extrinsic aging stimuli^2^. iii) Biomechanical properties are characterized and quantified using inconsistent and often insufficient metrics. iv) Many variables impact skin biomechanics that may not be considered among studies, e.g., degree of hydration^33^. v) Finally, both genetic and behavioral diversity lead to large inter-subject variability^34,35^.

Thus, despite our obvious interest, human skin studies have so far failed to paint a clear picture of the age- and sex-dependent mechanical properties of skin. Others have therefore resorted to studying skin in animals where genetic homogeneity, control of environmental factors, and tissue availability allow for more rigorous studies of skin mechanics. As in most other scientific disciplines, the use of rats and mice far exceeds the use of other animal models^36,37^. The popularity of murine models stems from their low cost, easy handling and housing, genetic malleability, and genetic similarity to humans^4,38^. Unfortunately, even with the use of murine models, many questions about age- and sex-dependent skin mechanics remain. Of the prior studies on the effect of age on murine skin, none have included sex as a variable, few have included very old animals, and even fewer have combined *in-vivo* mimicking test modes with compositional, structural, and microstructural investigations.

We set out to fill the significant gaps in our knowledge about how age and sex, and their interactions, affect skin mechanics. Given the challenges with human studies and prior test modalities, we investigate the age-sex interactions in mouse skin using mechanical tests that mimic the deformation that skin experiences in-vivo. Namely, we test skin mechanics under biaxial tension. In addition, we correlate potential differences between young and old mice as well as between male and female mice to compositional, structural, and microstructural differences, which we determine through a combination of collagen assays, histology, and two-photon microscopy. Thereby, our study provides insight into the remaining fundamental questions about how skin mechanics depend on age and sex.

## RESULTS

### Female skin is more prestretched than male skin

Prestrain quantifies the amount by which skin expands (positive values) or contracts (negative values) after being excised. **Figure 1** compares our prestrain measurements in young and old mice, male and female mice, between the dorsal and ventral sides, and between the lateral and the cranial-caudal directions. In general, we found that mouse skin contracts when excised, i.e., skin is stretched in-situ. Moreover, we found that prestrain differs significantly between the lateral and the cranial-caudal directions (p<0.001) and between the dorsal and ventral sides (p<0.0001). Most interestingly, we found that prestrain is sex dependent (p<0.001) with female skin contracting more than male mice. However, no age dependence of prestrain was detected (p=0.835).

**Figure 1.**
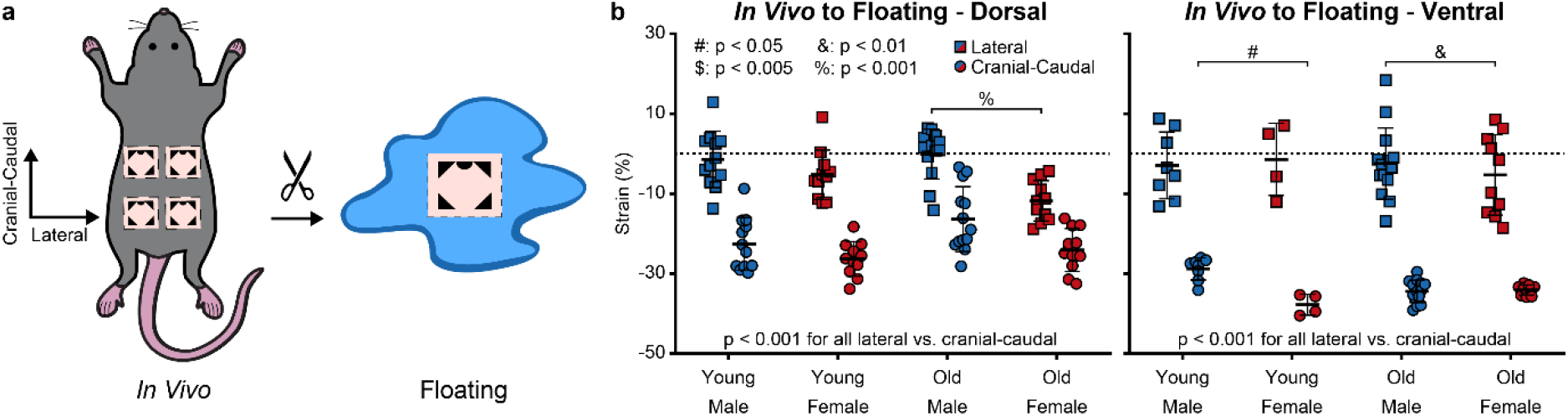
Female skin is more prestretched than male skin. (**a**) Skin samples were stamped in-situ and then excised. Based on the deformation of the skin stamp, we computed prestrain as a measure of the degree to which skin expanded (positive values) or contracted (negative values) after excision. (**b**) Comparison of skin prestrain between young and old mice, male and female mice, the dorsal and ventral sides, and directions. Young Male Dorsal n=12, Young Male Ventral n=8, Young Female Dorsal n=11, Young Female Ventral n=4, Old Male Dorsal n=13, Old Male Ventral n=14, Old Female Dorsal n=11, Old Female Ventral n=10.

### Old skin is less stiff than young skin and old female skin is relatively stiffer than old male skin

The mechanical behavior of skin is nonlinear and yields a convex stress-stretch curve, see **Supplementary Figure 1**. To statistically compare these nonlinear curves between groups, we identified three characteristic parameters: i) the slope of the stress-stretch curve at small stretches, which we call the “toe stiffness”, ii) the slope of the stress-stretch curve at large stretches, which we call “calf stiffness”, and iii) the stretch at 150 kPa, chosen to represent the stretch at which the toe region transitions to the calf region, see **Supplementary Figure 2**. Comparing these measures of skin biomechanics in **Figure 2a**, we found that the toe stiffness is well preserved across all our groups with differences only between the lateral and the cranial-caudal direction (p<0.001), but with no differences between young and old (p=0.705), male and female (p=0.621), or between the dorsal and ventral sides (p=0.171). Specifically, we found that the toe stiffness is larger in the lateral direction than in the cranial-caudal direction. In contrast, while the calf stiffness also differed significantly between the lateral and the cranial-caudal direction (p<0.0001) – with calf stiffness being higher in the latter direction – it also differed with age (p<0.0001), see **Figure 2b**. That is, young skin had a higher calf stiffness than old skin. Interestingly, while sex did not differ as a main effect (p=0.676), we did find a significant interaction between age and sex (p=0.047). Thus, it appears age-related reduction in skin stiffness was less in female skin than male skin, i.e., female skin became relatively stiffer when compared to male skin. No difference in calf stiffness was found between the dorsal and the ventral side (p=0.483). Finally, we compared stretches at 150 kPa which essentially measures how extendable samples are before stiffening, see **Figure 2c**. Here we found that samples are more extendable in the cranial-caudal direction than in the lateral direction (p<0.0001). No differences with age, sex, or dorsal/ventral side were significant (p=0.140, p=0.084 and, p=0.734, respectively).

**Figure 2.**
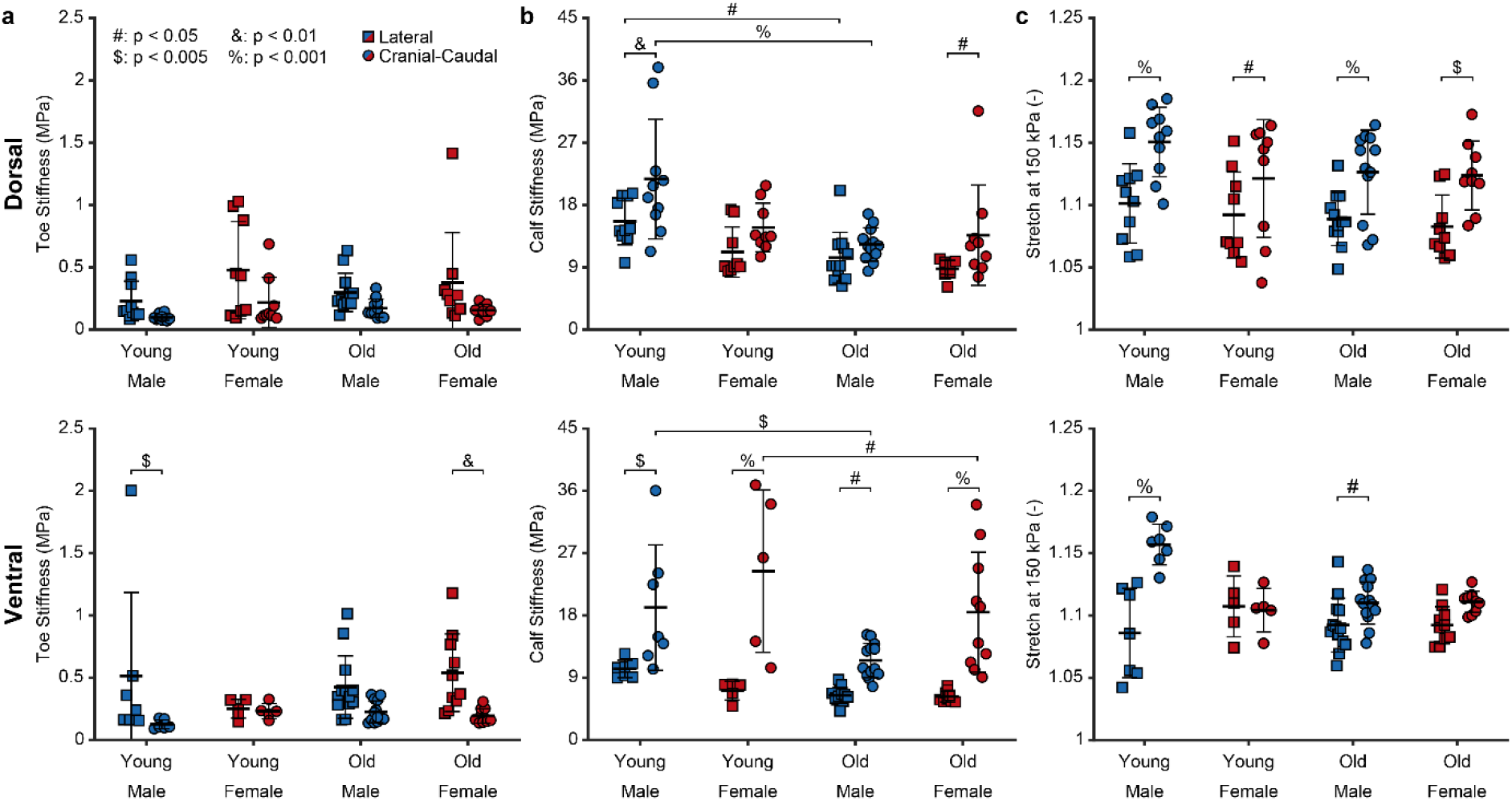
Old skin is less stiff than young skin and old female skin is relatively stiffer than old male skin. (**a**) Comparison of toe stiffness of dorsal and ventral skin samples between young and old mice, male and female mice, and directions. (**b**) Comparison of calf stiffness of dorsal and ventral skin samples between young and old mice, male and female mice, and directions. (**c**) Comparison of stretch at 150 kPa of dorsal and ventral skin samples between young and old mice, male and female mice, and directions. Young Male Dorsal n=10, Young Male Ventral n=7, Young Female Dorsal n=9, Young Female Ventral n=5, Old Male Dorsal n=12, Old Male Ventral n=13, Old Female Dorsal n=9, Old Female Ventral n=10.

### Old and female skin has more hypodermal fat, a thinner dermis, and less collagen

**Figure 3** compares the structure and composition of our skin samples. In these data we found three clear trends related to age and sex. First, age reduced dermal thickness (p<0.0001) and so did female sex (p<0.0001). Second, age increased hypodermal thickness (p=0.007) and so did female sex (p<0.001). Third, age reduced collagen concentration as measured by hydroxyproline quantification (p=0.014) and so did female sex (p=0.001). In addition, we found that hypodermal and muscular thickness differed with dorsal and ventral side (p<0.001 and p=0.005, respectively), but not epidermal thickness, dermal thickness, or collagen density (p=0.077, p=0.115 and p=0.994, respectively).

### Skin’s microstructural organization is preserved across age and sex

Finally, **Figure 4** compares the samples’ microstructural organization via depth-dependent collagen orientation probability maps. **Figure 4a** illustrates our visualization technique where we establish orientation probability functions for each imaging depth and project those onto a 2D plane. **Figure 4b** shows the orientation probability maps as a function of imaging depth. These maps show clear trends that are preserved across age, sex, and side. Specifically, Second Harmonic Generation (SHG)-derived collagen orientation is the most probable at 90° as measured against the cranial-caudal direction, i.e., in lateral direction. This means direction is preserved through the tissue depth. However, the fiber dispersion increases with depth (p<0.0001). In other words, while the mean fiber direction remains at 90°, the probability of fibers to deviate from this direction increases at deeper skin levels. Neither the mean fiber orientation nor the fiber dispersion appeared to differ as a function of age, sex, or side, i.e., dorsal versus ventral.

**Figure 3.**
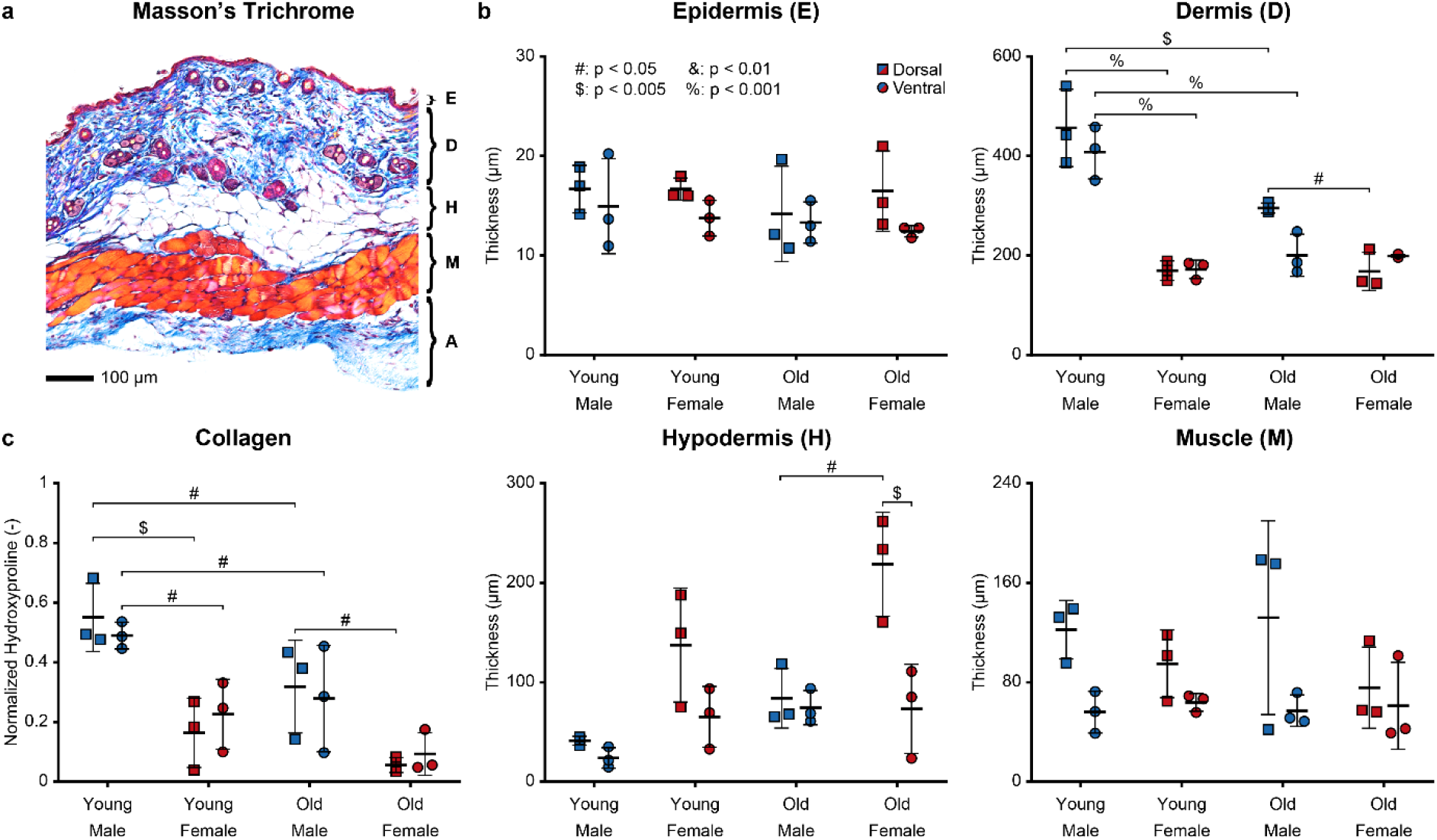
Old and female skin has more hypodermal fat, a thinner dermis, and less collagen. (**a**) Exemplary Masson’s Trichrome stain of mouse skin with layer definitions: epidermis (E), dermis (D), hypodermis (H), muscle (M) and adventitia (A). (**b**) Comparison of layer thickness between skin from young and old mice, male and female mice, and the dorsal and ventral sides. (**c**) Comparison of hydroxyproline content (normalized by total protein content) between skin from young and old mice, male and female mice as well as the dorsal and ventral sides. n=3 per group.

**Figure 4.**
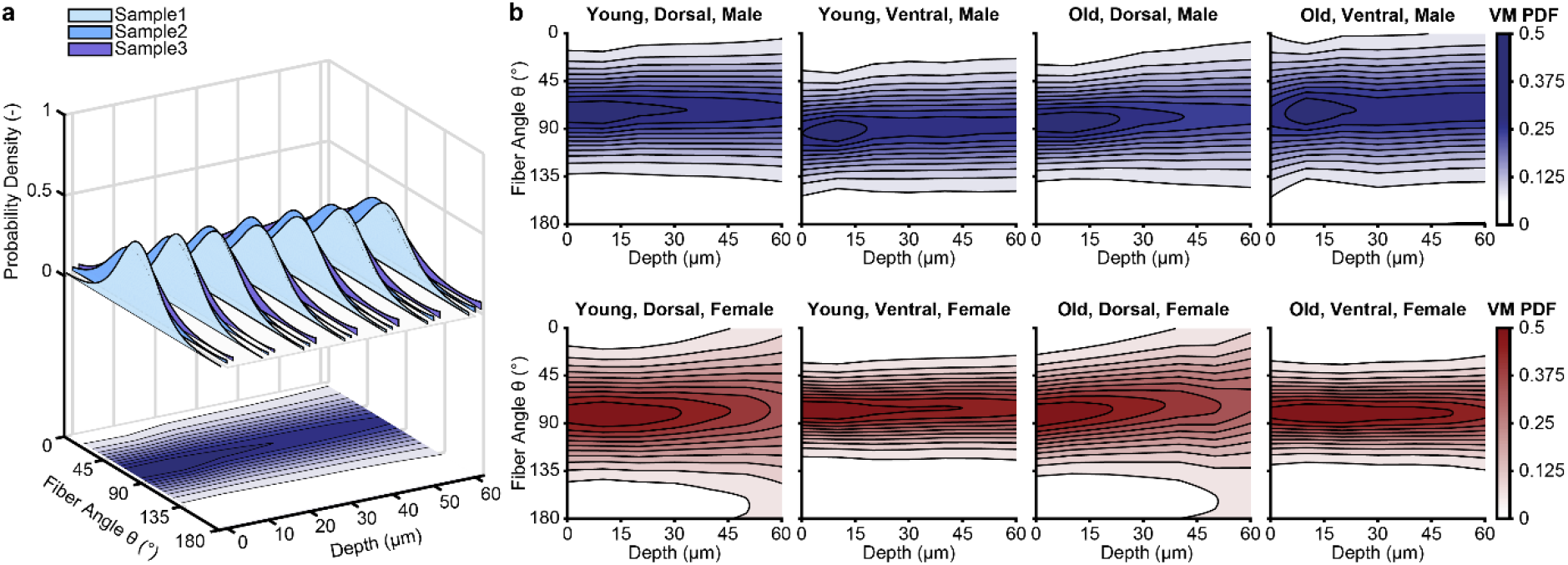
Microstructural organization is preserved across age and sex. (**a**) Depiction of our projection method with which we visualize depth-dependent 3D distributions in panel (**b**). (**b**) Comparison of skin’s depth-dependent fiber orientation probability between young and old, male and female skin, and the dorsal and ventral sides. n=3 per subpanel.

## DISCUSSION

Our goal was to answer fundamental questions about how skin mechanics depend on age and sex. To this end, we used young and old, male and female mice and tested their skin through a combination of biaxial testing, histology, collagen assays, and two-photon microcopy. In total we tested 156 skin samples from 52 mice spanning both sexes and an age range from 12 weeks to 80 weeks, which roughly represents mature adulthood (∼20-30 years of human age) and old age (∼55-70 years of human age)^37^. The most critical lesson we learned from these experiments is that the mechanics of mouse skin change with age and differ between male and female mice. Moreover, we learned that some sex differences in skin stiffness are age induced.

Specifically, we learned that age reduces skin calf stiffness but not toe stiffness. In other words, at small strains the mechanics of young and old skin do not differ, but at large strains they do. Given that collagen is attributed with dominating the skin mechanics at large strains, while elastin is argued to dominate skin mechanics at small strains, this age-induced difference in calf stiffness implicates collagen in this aging mechanism^39,40^. Indeed, our data also show that skin collagen density declines with age. Please note that stiffness is a normalized quantity and is independent of sample thickness. Therefore, our finding that dermal thickness also decreased with age compounds our finding on stiffness and means that skin’s structural stiffness (which accounts for skin thickness^41^) is reduced even more with age. Interestingly, our data on age-induced changes in skin stiffness in mice directly contradicts those findings by Lynch et al. who found that “tangent stiffness” – akin to our “calf stiffness” – increases in mice of similar age to ours^42^. Importantly, they used uniaxial tensile testing rather than biaxial testing as we did. We suspect that our use of a biaxial testing mode reveals age-induced changes in cross-fiber interactions that led to different findings than when using a uniaxial testing mode where cross-fiber interactions play less of a role. Our findings also disagree with our own prior work where we did not observe age-induced changes in mouse skin mechanics^43^. It should be noted, however, that mice in our prior study were roughly half the age of our current study and that we chose the age of mice in our current study specifically to overcome the limitation of our prior work. In contrast, our findings that mouse skin thickness and collagen content decreased with age is well supported by others in mice^44^, including Lynch et al^42^.

As for sex differences, we found that neither toe stiffness nor calf stiffness in young mice varied between female and male skin. This is despite significantly varied collagen densities. One possible explanation for this phenomenon lies in stiffening mechanisms other than collagen density, such as collagen structure, type, or degree of crosslinking^44^. Please note, however, that dermal thickness varied significantly between young male and female skin. Thus, structural stiffness between sexes differed with young male skin being structurally stiffer than young female skin (roughly by an order of two based on their relative dermal thickness). Notably, despite no differences in stiffness between young male and female skin, stiffness did differ between old male and female skin, with old female mice having significantly stiffer skin than old male mice. That is, age induced a sex difference in skin stiffness. This age-induced relative stiffening in female skin was likely driven by a comparatively smaller decline in collagen density in female skin than in male skin, as we showed in our work. Additionally, or alternatively, differences in age-mediated changes in skin stiffness could also stem from mice’s sex-dependent response to hormonal changes^45^. It should also be mentioned that none of the above differences stem from changes in skin microstructure, which did not differ with age or sex.

We also observed that female skin is more prestretched than male skin independent of age. This is interesting considering skin’s nonlinear stress-stretch relationship where prestretch leads to a shift of skin’s “operating range” toward higher stretches and thus a stiffer portion of the stress-stretch curve. In other words, the higher prestretch in female skin could be a compensatory mechanism to make up for its reduced structural stiffness when compared to male skin^46^. Unfortunately, we found no other prior studies on sex-dependent differences in mouse skin mechanics that we could compare our findings to; let alone studies that study sex and age interaction.

In addition to above novel findings on age- and sex-dependent mechanics of skin, we also confirmed prior findings. For example, we reconfirmed that skin mechanics differ directionally, i.e., skin is anisotropic, and that differences exist between locations, i.e., skin is also heterogeneous. Those findings agree well with our and others’ findings in murine skin^39,40,43,47–49^.

Of course, the ultimate goal of our and others’ work on mice is to translate our findings to humans. Toward this goal, a first critical step is to compare our findings to those on human skin. Before doing so, we’d like to raise a word of caution. Studies on human skin have employed a plethora of methodologies^50^. Here we only focus on comparing our results to those studies that used similar methodologies to ours (i.e., mechanical testing via uniaxial or biaxial testing) and where measures of skin mechanics were clearly defined. In contrast, we ignore those methods that make use of torsion or suction devices (e.g., cutometers^19^ and similar apparatuses), which we argue likely test not only the mechanics of skin but also the mechanics of the underlying subcutaneous tissues. This effectively excludes *in-vivo* studies and focuses on findings from post-mortem *in-vitro* studies.

Interestingly, most *in-vitro* studies of skin mechanics date back to the last century, while more recent work has primarily focused on *in-vivo* measurements. Among those early works, there is strong agreement that the mechanics of human skin changes with age^3,27–29^. Here we focus on prior reports of elastic modulus and extensibility, which are most comparable to our measures of calf stiffness and stretch at 150 kPa. For example, Holzmann et al. found that the stiffness and extensibility of adult skin decreased with age^27^. This finding is supported by Vogel, who also found that stiffness and extensibility of adult skin decreased with age^28^. Daly et al. found that extensibility at low strains decreased with age but did not find a difference in stiffness with age^3^. A very recent study compared the mechanical properties of human skin by age and location and found that the stiffness of skin from some body parts decreased with age, while the stiffness of skin samples from other body parts showed no age effect. Thus, they found significant heterogeneity in skin mechanics and skin’s age dependence^30^. Based on this selected work, it appears that the stiffness and extensibility of human skin decrease with age, but also shows some disagreements likely as a function of different test methods and differences in sample origin^30,51^.

Together, work on human skin supports our findings in mice on decreasing calf stiffness with age, but disagrees with our negative finding on extensibility, i.e., we did not find that stretch at 150 kPa significantly differed as a function of age. This may point to physiological differences between mouse and human skin or may be due to differences in our measures of extensibility compared to others. Most surprising to us was that prior work overwhelmingly agreed that sex does not affect stiffness or extensibility of skin. For example, Holzmann et al. saw no sex difference in skin stiffness^27^. Similarly, Jansen et al did not see a sex difference in skin stiffness^29^, nor did Zwirner^30^. Thus, our findings on the dependence of mouse skin mechanics on sex apparently do not represent findings in human skin well.

As for skin thickness and collagen content: there is mostly agreement in prior work that the thickness of adult skin decreased with age^20,27,31,52^, albeit there are also some that found no differences or found that skin increased in thickness, especially where UV exposure was more likely^15^. Similarly, prior work noted an age-induced collagen decrease in adult skin^28^, while some work found no changes with age^17,18^. However, most studies on human skin agree that skin thickness and collagen density decrease with age. Thus, our findings on the age dependence of mouse skin thickness and collagen content represent those findings in human skin well. Also, most prior work found that human skin thickness does depend on sex with female skin being thinner than male skin^52^, just as we found in our work.

## CONCLUSION

In conclusion, we found that skin stiffness, thickness, and collagen content all decreased with age in mice. We also found that these changes were sex dependent. The change in stiffness was age induced, meaning that only in old mice, female skin stiffness was larger than male skin stiffness. Overall, our findings agree well with findings from *in-vitro* studies on human skin. A notable difference is our finding on sex differences in stiffness, which have not been observed in human skin.

## MATERIALS & METHODS

### Sample Preparation

To study and understand the age and sex differences of mouse skin, we used 12-week (young) and 80-week (old) C57BL/6 male and female mice. We strictly adhered to NIH’s Guide for Care and Use of Laboratory Animals and all animal procedures described here were approved by the Institutional Animal Care and Use Committee at the University of Texas at Austin under #AUP-2020-00054. Following the humane sacrifice of the mice via CO_2_ inhalation, we removed the hair from dorsal and ventral skin regions using clippers and a chemical depilatory agent (Nair, Church & Dwight Co., Inc., Ewing, NJ, USA). Next, we applied an ink stamp of known dimensions (6 mm x 6 mm square) to four dorsal and two ventral skin regions. Next, we excised those stamped skin regions to obtain 12 mm x 12 mm square skin samples, for a total of six skin samples per mouse. We allocated all samples into four separate groups for mechanical testing, histology, compositional assays, and two-photon microscopy.

### Prestrain Calculation

Upon excision, we floated the skin samples from each group (young/old, male/female and dorsal/ventral) on a layer of 1x PBS at room temperature with the epidermis facing up. In this approximately stress-free, floating configuration, we photographed the specimens with their stamped profile clearly visible on a calibrated grid. After taking these images, we stored the samples at 4 °C in 1x PBS in preparation for subsequent biomechanical testing, see next paragraph. To compute prestrain, we identified the coordinates of the stamp’s corners from above photographs in a custom MATLAB (MATLAB R2020b, Mathworks, Natick, MA) program. Next, we bilinearly interpolated the deformation between the *in vivo* configuration, i.e., the original 6 × 6 mm square, and this floating configuration. Based on the resulting deformation field ***φ***_*p*_ we computed the deformation gradient tensor ***F***_*p*_ as the material gradient between both configurations, i.e., ***F***_*p*_ = ∇_*X*_***φ***_*p*_. Then, we quantified prestrain in terms of the Green-Lagrange strain tensor, 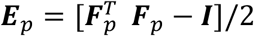, where ***I*** is the second order identity tensor. Note, if skin expanded after excision, Lagrangian strains were positive, and if skin contracted, these strains were negative.

### Biaxial Testing

Prior to mounting the samples for biaxial testing, we measured sample thickness at four locations using a digital thickness gauge (547-500S, Mitutoyo Corp., Kawasaki, Japan). Then we speckled the epidermal side with graphite powder. After mounting the sample on our biaxial device (Biotester, Cellscale, Waterloo, ON, Canada) and submerging the sample in 37 °C 1x PBS, we preloaded the tissue equibiaxially (the same in all directions) to 50 mN to establish a consistent reference state and to remove tissue slack. Next, we performed 20 preconditioning cycles equibiaxially to 1000 mN and then conducted two final equibiaxial cycles to 1000 mN (force-controlled test at a quasi-static rake displacement rate of approximately 0.25 mm/s). We chose this force to induce large deformation in the skin samples without causing damage. While testing, we continuously captured images of the speckle pattern at 5 Hz for off-line digital image correlation. We performed all tests within 4 hours of excision. We acquired actual tissue strain during biaxial testing via digital image correlation of the recorded graphite pattern using the Cellscale image analysis software Labjoy. We chose the end of the second down stroke of the equibiaxial cycles as the reference configuration to yield the elastic deformation map and deformation gradient, i.e., ***F***^*e*^ = ∇_*X*_ ***φ***^*e*^. In the following analysis, we report stress-stretch data relative to the preloaded *in vitro* configuration, i.e., the mounted configuration after 50 mN of preload was applied. Moreover, we transformed load data into Cauchy stress via measurements of the sample’s thickness and width, which we projected into the current configuration under the assumption of tissue incompressibility. Please note that we excluded samples that were not loaded to at least 185 kPa.

### Histology

Upon excision, we immediately fixed those skin samples that were allocated for histology in 10% neutral buffered formalin for 24 hours, then transferred them directly to 70% ethanol. A commercial histology service (Histoserv Inc., Germantown, MD, USA) prepared all histological slides by embedding them in paraffin, sectioning them transversely to a thickness of 5 μm and staining them with Masson’s Trichrome. We subsequently acquired histological images on an upright microscope (BX53 Upright Microscope, Olympus, Tokyo, Japan) at 10x magnification. To measure layer thicknesses, we used a custom MATLAB program to load images of the Masson Trichrome stains. Therein, we manually measured the thickness of the epidermal, dermal, subcutaneous, muscular, and adventitial layers while blinded to the experimental group. We did so at 24 points along the length of the section before averaging those values for each sample.

### Quantitative Collagen Assay

Upon excision, we cryogenically stored the skin samples from each group at −80 °C in a 9:1 ratio of DMEM:DMSO with protease inhibitor (ThermoFisher, A32953, Weltham, MA) until assayed. Immediately before testing, we rapidly thawed the samples to room temperature and acquired the wet mass of each sample. For every 10 mg of wet tissue mass, we added 100 mL DI water for homogenization. We hydrolyzed 100 mL of homogenate in 100 mL of 10 N NaOH at 120 °C for 1 h, after which we neutralized by adding 100 mL of 10 N HCl. After vortex mixing at 2000x g for 5 min, we transferred 10 mL of hydrolysate to each well in triplicate, which we allowed to evaporate to dryness on a 65 °C heating plate. We then followed the protocol provided with the Total Collagen Assay Kit (BioVision Inc, K406, Milpitas, CA, USA) and measured the colorimetric absorbance at 560 nm with a spectrophotometer (Tecan, Infinite 200 Pro, Männedorf, Switzerland) which we interpolated from a standard type-I linear fit curve (R^2^ = 0.99 ± 0.01). Additionally, we determined the total protein concentration of each sample using a Pierce™ Microplate BCA Protein Assay Kit (Thermo Fisher Scientific, 23252, Waltham, MA, USA) to calculate the normalized collagen content.

### Two-Photon Microscopy

Upon excision, we cryogenically stored the skin samples from each group at −80 °C in a 9:1 ratio of DMEM:DMSO with protease inhibitor. Immediately before imaging, we rapidly thawed the samples to room temperature and washed them with 1X PBS. We imaged the skin samples from each group under a two-photon microscope (Ultima IV, Bruker, Billerica, MA, USA) for the *in-vitro* collagen fiber orientation analysis via Second Harmonic Generation (SHG). We acquired all images epidermis up using a 20x water immersion objective (XLUMPLFLN, Olympus, Center Valley, PA, USA) at an excitation wavelength of 900 nm. We epi-collected the backscattered SHG through a PMT channel filter (460 ± 25 nm) and acquired a z-stack of images with a step size of 10 μm until the SHG intensity diminished (∼100 μm) at four different locations in the center of the tissue. To analyze the SHG images, we used the orientation distribution analysis with a Gaussian gradient method in ImageJ-FIJI OrientationJ (National Institutes of Health, Bethesda, MD, USA)^53^. Subsequently, we fit a symmetric von Mises distribution to the raw data to estimate the distribution’s location parameter 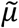 and localization parameter 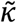 as a function of imaging depth^54^.

### Statistics

We conducted the statistical analyses in R (Version 4.1.2) where statistical significance was defined as p < 0.05. For all data except the thickness measurement of skin layers, we used a linear mixed model as implemented in the R package *afex*, while for thicknesses of skin layers, we performed a three-way ANOVA. All post-hoc analyses were conducted using Tukey tests. Where applicable, we reported data as mean with the standard deviation of the mean.

## DATA AVAILABILITY STATEMENT

All data is available through our Data Repository: https://dataverse.tdl.org/dataverse/age_sex_mouse_skin_mechanics.

## ACKNOWLEDGEMENTS

This work was partially supported by NSF grants to MKR (1916663, 2046148, 2105175, 2127925) and ABT (1916668), and by an NIH predoctoral fellowship to WDM (F31HL145976). The opinions, findings, and conclusions, or recommendations expressed are those of the authors and do not necessarily reflect the views of the National Science Foundation.

## AUTHOR CONTRIBUTIONS

Conceptualization: GPS, WDM, ABT, MKR; Data Curation: C-YL; Formal Analysis: C-YL, GPS, SK; Investigation: C-YL, GPS; Software: WDM, SK; Supervision: WDM, ABT, MKR; Visualization: C-YL; Writing – Original Draft Preparation: C-YL, ABT, MKR; Writing – Review and Editing: C-YL, GPS, SK, WDM, ABT, MKR; Funding Acquisition: ABT, MKR.

## SUPPLEMENTARY FIGURES

**Supplementary Figure 1.**
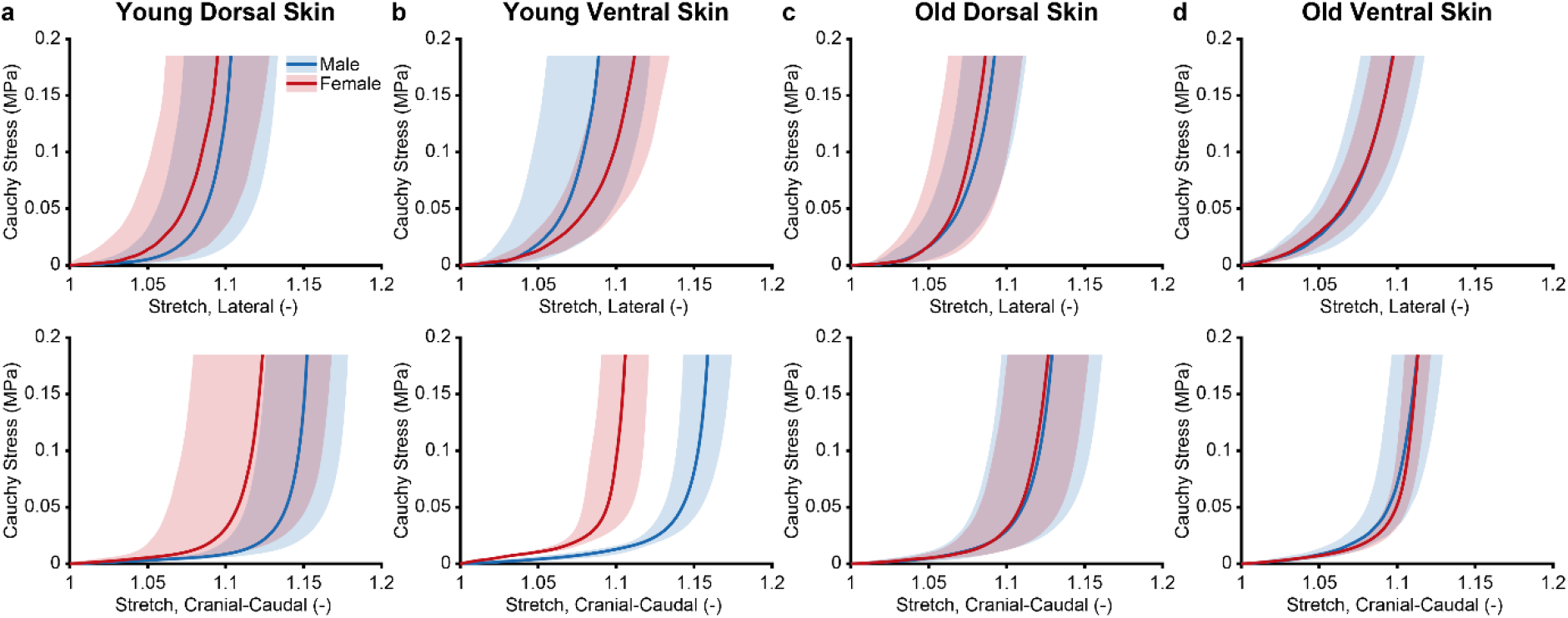
Mouse skin shows characteristic J-shaped stress-stretch relationship. Skin’s stress-stretch average curves (solid) with standard deviation (shaded) in young and old mice, male and female mice, the dorsal and ventral sides, and directions.

**Supplementary Figure 2.**
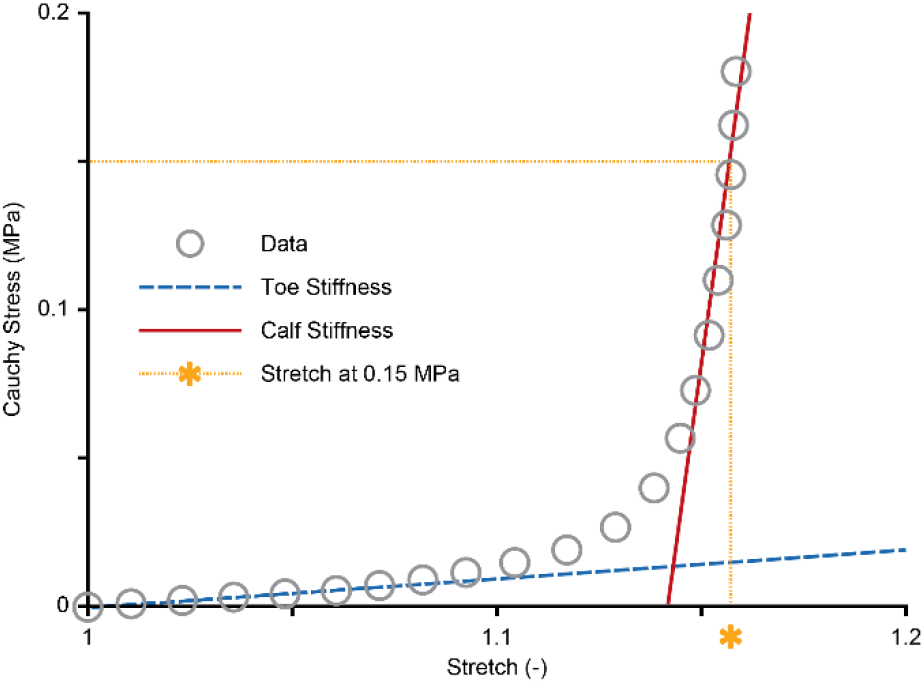
Depiction of mechanical metrics. Showing toe stiffness as the slope of the stress-stretch curve at small stretches, calf stiffness as the slope of the stress-stretch curve at large stretches, and the stretch at 150 kPa (i.e., 0.15 MPa).

